# Analytic Pearson residuals for normalization of single-cell RNA-seq UMI data

**DOI:** 10.1101/2020.12.01.405886

**Authors:** Jan Lause, Philipp Berens, Dmitry Kobak

## Abstract

**Background:** Standard preprocessing of single-cell RNA-seq UMI data includes normalization by sequencing depth to remove this technical variability, and nonlinear transformation to stabilize the variance across genes with different expression levels. Instead, two recent papers propose to use statistical count models for these tasks: Hafemeister & Satija [1] recommend using Pearson residuals from negative binomial regression, while Townes et al. [2] recommend fitting a generalized PCA model. Here, we investigate the connection between these approaches theoretically and empirically, and compare their effects on downstream processing.

**Results:** We show that the model of Hafemeister and Satija produces noisy parameter estimates because it is overspecified, which is why the original paper employs post-hoc smoothing. When specified more parsimoniously, it has a simple analytic solution equivalent to the rank-one Poisson GLM-PCA of Townes et al. Further, our analysis indicates that per-gene overdispersion estimates in Hafemeister and Satija are biased, and that the data are in fact consistent with the overdispersion parameter being independent of gene expression. We then use negative control data without biological variability to estimate the technical overdispersion of UMI counts, and find that across several different experimental protocols, the data are close to Poisson and suggest very moderate overdispersion. Finally, we perform a benchmark to compare the performance of Pearson residuals, variance-stabilizing transformations, and GLM-PCA on scRNA-seq datasets with known ground truth.

**Conclusions:** We demonstrate that analytic Pearson residuals strongly outperform other methods for identifying biologically variable genes, and capture more of the biologically meaningful variation when used for dimensionality reduction.

## 1 Introduction

The standard preprocessing pipeline for single-cell RNA-seq data includes sequencing depth normalization followed by log-transformation [3, 4]. The normalization aims to remove technical variability associated with cell-to-cell differences in sequencing depth, whereas the log-transformation is supposed to make the variance of gene counts approximately independent of the mean expression. Two recent papers argue that neither step works very well in practice [1, 2]. Instead, both papers suggest to model UMI (unique molecular identifier) data with count models, explicitly accounting for the cell-to-cell variation in sequencing depth (defined here as the total UMI count per cell). Hafemeister & Satija [1] use a negative binomial (NB) regression model (`scTransform` package in R), while Townes et al. [2] propose Poisson generalized principal component analysis (GLM-PCA). These two models are seemingly very different.

Here we show that the model used by Hafemeister & Satija [1] has a too flexible parametrization, resulting in noisy parameter estimates. As a consequence, the original paper employs post-hoc smoothing to correct for that. We show that a more parsimonious model produces stable estimates even without smoothing and is equivalent to a special case of GLM-PCA. We then demonstrate that the estimates of gene-specific overdispersion in the original paper are strongly biased, and further argue that UMI data do not require gene-specific overdispersion parameters to account for technical noise. Rather, the technical variability is consistent with the same overdispersion parameter shared between all genes. We use available negative control datasets to estimate this technical overdispersion. Furthermore, we compare Pearson residuals, GLM-PCA, and variance-stabilizing transformations for highly variable gene selection and as data transformation for downstream processing.

Our code in Python is available at http://github.com/berenslab/umi-normalization. Analytic Pearson residuals will be included into upcoming `Scanpy` 1.9 [5].

## 2 Results

### 2.1 Analytic Pearson residuals

A common modeling assumption for UMI or read count data without biological variability is that each gene *g* takes up a certain fraction *p*_*g*_ of the total amount *n*_*c*_ of counts in cell *c* [2, 6, 7,8,9,10]. The observed UMI counts *X*_*cg*_ are then modelled as Poisson or negative binomial (NB) [11] samples with expected value *μ*_*cg*_ = *p*_*g*_*n*_*c*_ without zero-inflation [10, 12]:

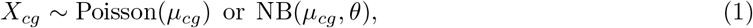

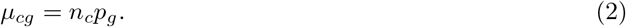

The Poisson model has a maximum likelihood solution (see Methods) that can be written in closed form as 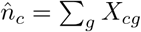 (sequencing depths), 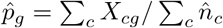, or, put together,

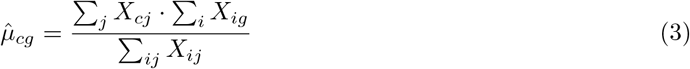

For the negative binomial model this holds only approximately. Using this solution, the Pearson residuals are given by

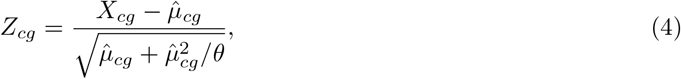

where 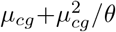 is the NB variance and *θ* → ∞ gives the Poisson limit. The variance of Pearson residuals is, up to a constant, equal to the Pearson *χ*^2^ goodness-of-fit statistic [13] and quantifies how much each gene deviates from this constant-expression model. As pointed out by Aedin Culhane [14], singular value decomposition of the Pearson residuals under the Poisson model is known as *correspondence analysis* [15, 16, 17, 18], a method with a longstanding history [19].

Hafemeister & Satija [1] suggested using Pearson residuals from a related NB regression model for highly variable gene (HVG) selection and also as a data transformation for downstream processing. In parallel, Townes et al. [2] suggested using deviance residuals (see Methods) from the same Poisson model as above for HVG selection and also for PCA as an approximation to their GLM-PCA. In the next sections we discuss the relationships between these approaches.

### 2.2 The regression model in scTransform is overspecified

Hafemeister & Satija [1] used the 33k PBMC (peripheral blood mononuclear cells, an immune cell class that features several distinct subpopulations) dataset from 10X Genomics in their work on normalization of UMI datasets. For each gene *g* in this dataset, the authors fit an independent NB regression

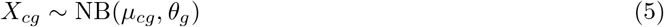

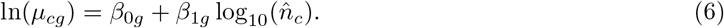

Here *θ*_*g*_ is the gene-specific overdispersion parameter, 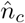 are observed sequencing depths as defined above, and *β*_0*g*_ and *β*_1*g*_ are the gene-specific intercept and slope. The natural logarithm follows from the logarithmic link function that is used in NB regression by default. The original paper estimates *β*_0*g*_ and *β*_1*g*_ using Poisson regression, and then uses the obtained estimates to find the maximum likelihood estimate of *θ*_*g*_. The resulting estimates for each gene are shown in Figure 1a−c, reproducing Figure 2A from the original paper.

**Figure 1:**
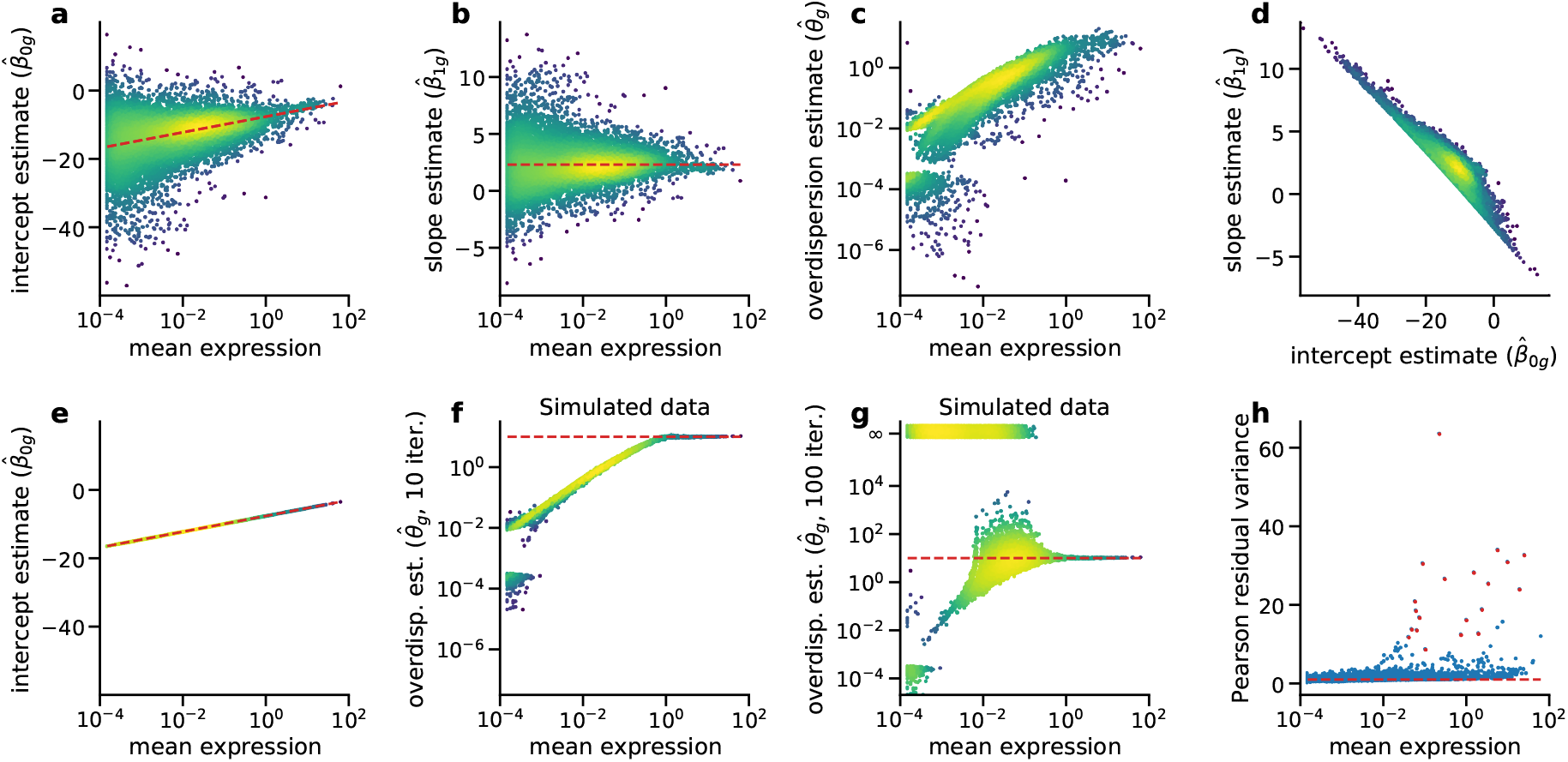
Regression model of Hafemeister & Satija [1] compared to the offset model. Each dot corresponds to a model fit to the counts of a single gene in the 33k PBMC dataset (10x Genomics, *n* = 33 148 cells). Following Hafemeister & Satija [1], we included only the 16 809 genes that were detected in at least five cells. Color denotes the local point density from low (blue) to high (yellow). Expression mean was computed as 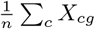. **a:** Intercept estimates 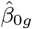 in the original regression model. Dashed line: Analytic solution for 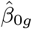 in the offset model we propose. **b:** Slope estimates 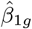 . Dashed line: *β*_1*g*_ = ln(10) *≈* 2.3. **c:** Overdispersion estimates 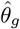. **d:** Relationship between slope and intercept estimates (*ρ* = *−*0.91). **e:** Intercept estimates in the offset model, where the slope coefficient is fixed to 1. Dashed line shows the analytic solution, which is a linear function of gene mean. **f:** Overdispersion estimates 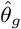 on simulated data with true *θ* = 10 (dashed line) for all genes. **g:** Overdispersion estimates 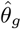 on the same simulated data as in panel (f), but now with 100 instead of 10 iterations in the theta.ml() optimizer (R, MASS package). Cases for which the optimization diverged to infinity or resulted in spuriously large estimates 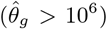 are shown at 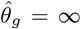 with some jitter. Dashed line: true value *θ* = 10. **h:** Variance of Pearson residuals in the offset model. The residuals were computed analytically, assuming *θ* = 100 for all genes. Following Hafemeister & Satija [1], we clipped the residuals to a maximum value of 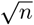. Dashed line indicates unit variance. Red dots show the genes identified in the original paper as most variable.

**Figure 2:**
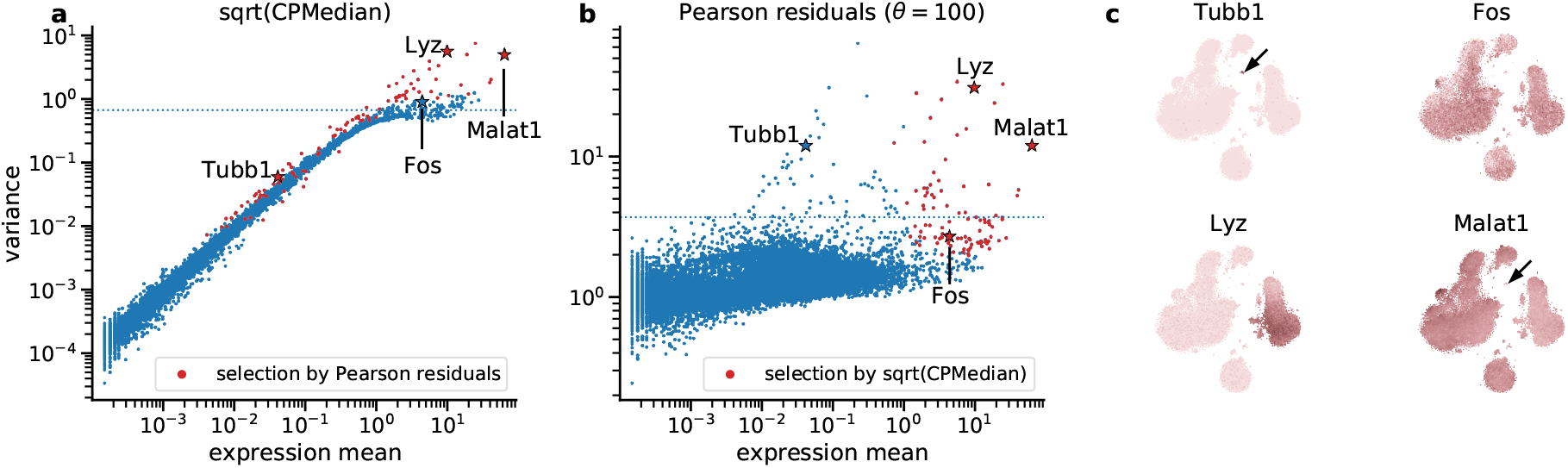
Selection of variable genes. In the first two panels, each dot shows the variance of a single gene in the PBMC dataset after applying a normalization method. The dotted horizontal line shows a threshold adjusted to select 100 most variable genes. Red dots mark 100 genes that are selected by the other method, i.e. that are above the threshold in the other panel. Stars indicate genes shown in the last panel. **a:** Gene variance after sequencing depth normalization, median-scaling, and the square-root transformation. **b:** Variance of Pearson residuals (assuming *θ* = 100). **c:** t-SNE of the entire PBMC dataset (see Additional File 1: Figure S4), colored by expression of four example genes (after sequencing depth normalization and square-root transform). Platelet marker *Tubb1* with low average expression is only selected by Pearson residuals. Arrows indicate the platelet cluster. *Fos* is only selected by the square root-based method, and does not show a clear type-specific expression pattern. *Malat1* (expressed everywhere apart from platelets) and monocyte marker *Lyz* with higher average expression are selected by both methods.

The authors observed that the estimates 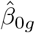 and 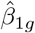 were unstable and showed high variance for genes with low average expression (Figure 1a−b). They addressed this with a ‘regularization’ procedure that re-set all estimates to the local kernel average estimate for a given expression level. This is similar to some approaches to bulk RNA-seq analysis [6, 7] but with post-hoc correction instead of Bayesian shrinkage. This kernel smoothing resulted in an approximately linear increase of the intercept with the logarithm of the average gene expression (Figure 1a) and an approximately constant slope value of 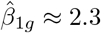 (Figure 1b). The nature of these dependencies was left unexplained. Moreover, we found that 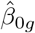 And 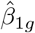 were strongly correlated (*ρ* = −0.91), especially for weakly expressed genes (Figure 1d).

Together, these clear symptoms of overfitting suggest that the regression model was overspecified.

Indeed, the theory calls for a less flexible model.As explained above, a common modeling assumption (Eq. 2) is that *μ*_*cg*_ = *p*_*g*_*n*_*c*_, or equivalently

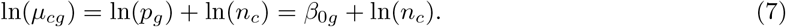

We see that under this assumption, the slope *β*_1*g*_ does not need to be fit at all and should be fixed to 1, if ln(*n*_*c*_) is used as predictor. Not only does this suggest an alternative, simpler parametrization of the model, but it also explains why Hafemeister & Satija [1] found that 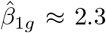: they used log_10_(*n*_*c*_) = ln(*n*_*c*_)*/* ln(10) instead of ln(*n*_*c*_) as predictor, and so obtained ln(10) *≈* 2.3 as the average slope.

Under the assumption of Eq. 7, a Poisson or NB regression model should be specified using ln(*n*_*c*_) as predictor with a fixed slope of 1, a so-called *offset* (Eqs. 5 and 7). This way, the resulting model has only one free parameter and is not overspecified. Moreover, the Poisson offset model is equivalent to Eqs. 1−2 and so, as explained above, has an analytic solution

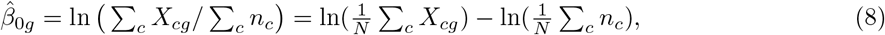

which forms a straight line when plotted against the log-transformed average gene expression 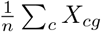 (Figure 1e). This provides an explanation for the linear trend in 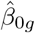 in the original two-parameter model (Figure 1a).

In practice, our one-parameter offset model and the original two-parameter model after smoothing arrive at qualitatively similar results (Figure 1h). However, we argue that the one-parameter model is more appealing from a theoretical perspective, has an analytic solution, and does not require post-hoc averaging of the coefficients across genes.

### 2.3 The offset regression model is equivalent to the rank-one GLM-PCA

The offset regression model turns out to be a special case of GLM-PCA [2]. There, the UMI counts are modeled as

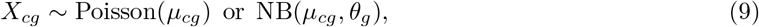

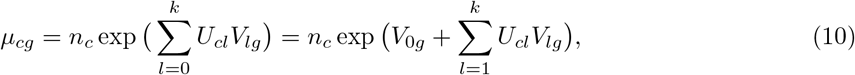

assuming *k* + 1 latent factors, with *U* and *V* playing the role of principal components and corresponding eigenvectors in standard PCA. Importantly, the first latent factor is constrained to *U*_*c*0_ = 1 for all cells *c*, such that *V*_0*g*_ can be interpreted as gene-specific intercepts. If the data are modeled without any further latent factors, Eq. 10 reduces to

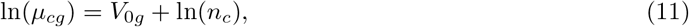

which is identical to Eq. 7 with *V*_0*g*_ = *β*_0*g*_. This shows that the proposed one-parameter offset regression model is exactly equivalent to the intercept-only rank-one GLM-PCA.

### 2.4 Overdispersion estimates in scTransform are biased

After discussing the overparametrization of the systematic component of the `scTransform` model, we now turn to the NB noise model employed by Hafemeister & Satija [1]. The 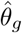 estimates in the original paper are monotonically increasing with the average gene expression, both before and after kernel smoothing (Figure 1c). This suggests that there is a biologically meaningful relationship between the expression strength and the overdispersion parameter *θ*_*g*_. However, this conclusion is in fact unsupported by the data.

To demonstrate this, we simulated a dataset with NB-distributed counts 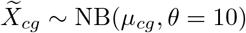 with *μ*_*cg*_ given by Eq. 3 using *X*_*cg*_ of the PBMC dataset. Applying the original estimation procedure to this simulated dataset showed the same positive correlation of 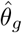 with the average expression as in real data (Figure 1f), strongly suggesting that it does not represent an underlying technical or biological cause, but only the estimation bias. Low-expressed genes had a larger bias and only for genes with the highest average expression was the true *θ* = 10 estimated correctly.

Moreover, the 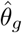 estimates strongly depended on the exact details of the estimation procedure. Using

the `theta.ml()` R function with its default 10 iterations, as Hafemeister & Satija [1] did, led to multiple convergence warnings for the simulated data in Figure 1f. Increasing this maximum number of iterations to 100 eliminated most convergence warnings, but caused 49.9% of the estimates to diverge to infinity or above 10^10^ (Figure 1g). These instabilities are likely due to shallow maxima in the NB likelihood w.r.t. *θ* [20].

The above arguments show that the overdispersion parameter estimates in Hafemeister & Satija [1] for genes with low expression were strongly biased. In practice, however, the predicted variance *μ* + *μ*^2^*/θ* is only weakly affected by the exact value of *θ* for low expression means *μ*, and so the bias reported here does not substantially affect the Pearson residuals (see below). Also, many of the weakly expressed genes may be filtered out during preprocessing in actual applications. We note that large errors in NB overdispersion parameter estimates have been extensively described in other fields, with simulation studies showing that estimation bias occurs especially for low NB means, small sample sizes, and large true values of *θ* [21, 22, 23], i.e. for samples that are close to the Poisson distribution. Note also that post-hoc smoothing [1] can reduce the variance of the 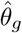 estimates, but does not reduce the bias.

### 2.5 Negative control datasets suggest low overdispersion

To avoid noisy and biased estimates, we suggest to use one common *θ* value shared between all genes. Of course, any given dataset would be better fit using gene-specific values *θ*_*g*_. However, our goal is not the best possible fit: We want the model to account only for technical variability, but not biological variability e.g. between cell types; this kind of variability should manifest itself as high residual variance.

Rather than estimating the *θ* value from a biologically heterogeneous dataset such as PBMC, we think it is more appropriate to estimate the technical overdispersion using negative control datasets, collected without any biological variability [12]. We analyzed several such datasets spanning different droplet- and plate-based sequencing protocols (10x Genomics, inDrop, MicrowellSeq) and compared the 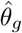 estimates to the estimates obtained using simulated NB data with various known values of *θ* ∈ {10, 100, 1000, ∞} For the simulations, we used the empirically observed sample sizes and sequencing depths. We found that across different protocols, negative control data were consistent with overdispersion *θ* ≈ 100 or larger (Additional File 1: Figure S1). The plateau at *θ* ≈ 10 in the PBMC data visible in Figure 1c could reflect biological and not technical variability. At the same time, negative control data did not exactly conform to the Poisson model (*θ* = ∞), but likely overdispersion parameter values (*θ* ≈ 100) were large enough to make the Poisson model acceptable in practice [10, 24, 25]. A Parallel work reached the same conclusion [26].

### 2.6 Analytic Pearson residuals select biologically relevant genes

Both Hafemeister & Satija [1] and Townes et al. [2] suggested to use Pearson/deviance residuals based on models that only account for technical variability, in order to identify biologically variable genes. Indeed, genes showing biological variability should have higher variance than predicted by such a model. As explained above, Pearson residuals in the model given by Eqs. 1−2 (or, equivalently, offset regression model or rank-one GLM-PCA) can be conveniently written in closed form:

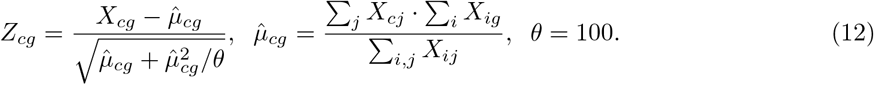

For most genes in the PBMC data, the variance of the Pearson residuals was close to 1, indicating that this model predicted the variance of the data correctly and suggesting that most genes did not show biological variability (Figures 1h). Using *θ* = 100 led to several high-expression genes selected as biologically variable that would not be selected with a lower *θ* (e.g. *Malat1*), but overall, using *θ* = 10, *θ* = 100, or even the Poisson model with *θ* =∞ led to only minor differences (Additional File 1: Figure S2). Using analytic Pearson residuals for HVG selection yielded a very similar result compared to using Pearson residuals from the smoothed regression presented in Hafemeister & Satija [1], with almost the same set of genes identified as biologically variable (Figure 1h, Additional File 1: Figure S2). This suggests that our model is sufficient to identify biologically relevant genes.

It is instructive to compare the variance of Pearson residuals to the variance that one gets after explicit sequencing depth normalization followed by a variance-stabilizing transformation. For Poisson data, the square root transformation 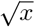 is approximately variance-stabilizing, and several modifications exist in the literature [27], such as the Anscombe transformation 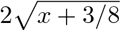 [28] and the Freeman-Tukey transformation 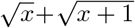 [29]. Normalizing UMI counts *X*_*cg*_ by sequencing depths *n*_*c*_ (and multiplying the result by the median sequencing depth ⟨*n*_*c*_⟩ across all cells; ‘median normalization’) followed by one of the square-root transformations has been advocated for UMI data processing [30, 31].

Comparing the gene variances after the square-root transformation (Figure 2a) with those of Pearson residuals (Figure 2b) in the PBMC dataset showed that the square-root transformation is not sufficient for variance stabilization. Particularly affected are low-expression genes that have variance close to zero after the square-root transform [32]. For example, platelet markers genes such as *Tubb1* have low average expression (because platelets are a rare population in the PBMC dataset) and do not show high variance after any kind of square-root transform (another example was given by the B-cell marker *Cd79a*). At the same time, Pearson residuals correctly indicate that these genes have high variance and are biologically meaningful (Figure 2c). For the genes with higher average expression, some differentially expressed genes like the monocyte marker *Lyz* or the above-mentioned *Malat1* showed high variance in both approaches. However, the selection based on the square-root transform also included high-expression genes like *Fos*, which showed noisy and biologically unspecific expression patterns (Figure 2c). Similar patterns were observed in the full-retina dataset [33] (Additional File 1: Figure S3).

The gene with the highest average expression in the PBMC dataset, *Malat1*, showed clear signs of biologically meaningful variability: e.g. it is not expressed in platelets (Figure 2c). While this gene is selected as biologically variable based on Pearson residuals with *θ* ≈ 100 as we propose (Figure 2b), it was not selected by Hafemeister & Satija [1] who effectively used *θ* ≈ 10 (Figure 1c,h, Additional File 1: Figure S2). This again suggests that *θ* ≈ 100 is more appropriate than *θ* ≈ 10 to model technical variability of UMI counts.

Pearson residuals may even be ‘too sensitive’ in that genes that are only expressed in a handful cells may get very large residual variance. Hafemeister & Satija [1] suggested clipping residuals to 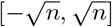. We found that this step avoids large residual variance in very weakly expressed genes (Additional File 1: Figure S2, see Methods for more details). The variance of unclipped Pearson residuals under the Poisson model (*θ* = ∞) was very similar to the Fano factor of counts after median normalization (Additional File 1: Figure S2) and less useful for HVG selection compared to the clipped residuals.

Lastly, gene selection by the widely-used log(1 + *x*)-transform as well as by the variance of deviance residuals as suggested by Townes et al. [2] led to very similar results as described above for the square-root transform: many biologically meaningful genes were not selected, as all three methods overly favored high-expression genes (Additional File 1: Figure S2). In conclusion, neither of these transformations is sufficiently variance-stabilizing. In practice, many existing HVG selection methods take the mean-variance relationship into account when performing the selection (e.g. `seurat` and `seurat v3` methods [34, 35] as implemented in `Scanpy` [5]). We benchmarked their performance in the next section.

### 2.7 Analytic Pearson residuals separate cell types better than other methods

Next, we studied the effect of different normalization approaches on PCA representations and t-SNE embeddings. The first approach is median normalization, followed by the square-root transform [30, 31]. We used 50 principal components of the resulting data matrix to construct a t-SNE embedding. The second approach is computing Pearson residuals according to Eq. 12 with *θ* = 100, followed by PCA reduction to 50 components. The third approach is computing 50 components of negative binomial GLM-PCA with *θ* = 100 [2]. We used the same initialization to construct all t-SNE embeddings to ease the visual comparison [38].

We applied these methods to the full PBMC dataset (Additional File 1: Figure S4), three retinal datasets [33, 36, 37] (Figure 3), and a large organogenesis dataset with *n* = 2 million cells [39] (Figure 4). For smaller datasets, the resulting embeddings were mostly similar, suggesting comparable performance between methods. Hafemeister & Satija [1] argued that using Pearson residuals reduces the amount of variance in the embedding explained by the sequencing depth variation, compared to sequencing depth normalization and log-transformation. We argue that this effect was mostly due to the large factor that the authors used for re-scaling the counts after normalization (Additional File 1: Figure S5): large scale factors and/or small pseudocounts (*ϵ* in log(*x* + *ϵ*)) are known to introduce spurious variation into the distribution of normalized counts [2, 40]. For the PBMC dataset, all three t-SNE embeddings showed similar amount of sequencing depth variation across the embedding space (Additional File 1: Figure S4g−i). Performing the embeddings on 1000 genes with the largest Pearson residual variance did not noticeably affect the embedding quality (Additional File 1: Figure S4).

**Figure 3:**
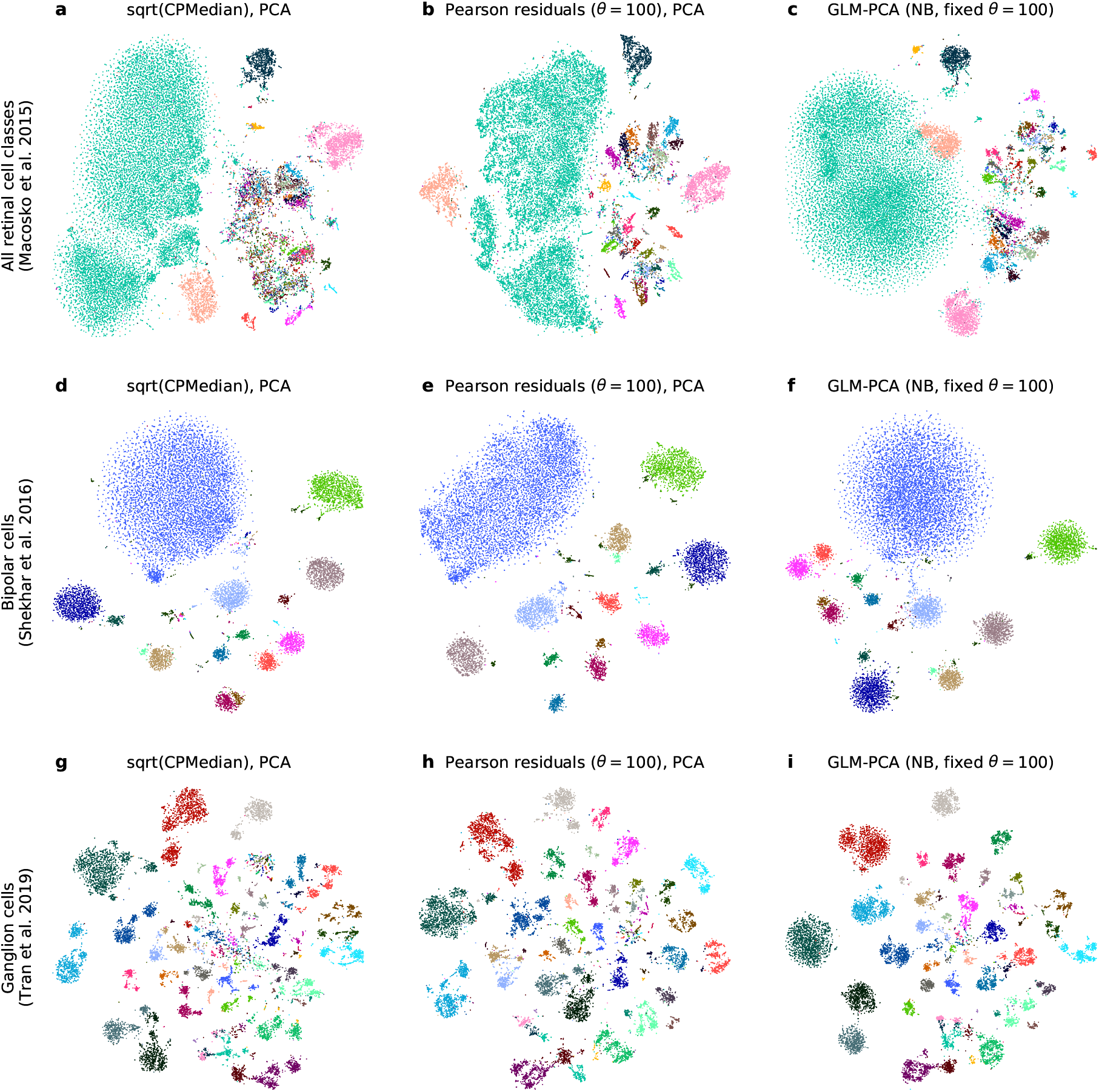
t-SNE embeddings of three retinal datasets. Panels in each column are based on a different data transformation method with PCA or GLM-PCA reduction to 50 dimensions (see Methods), and each row shows a different retinal dataset. We did not perform any gene selection here. Colors correspond to cell type labels provided by the original papers. **a−c:** Full-retina dataset (DropSeq) [33], containing all retinal cell types (including glia and vascular cells). 24 769 cells. **d−f:** Bipolar cell dataset (DropSeq) [36]. 13 987 cells. **g−i:** Retinal ganglion cell dataset (10X v2) [37]. 15 750 cells.

**Figure 4:**
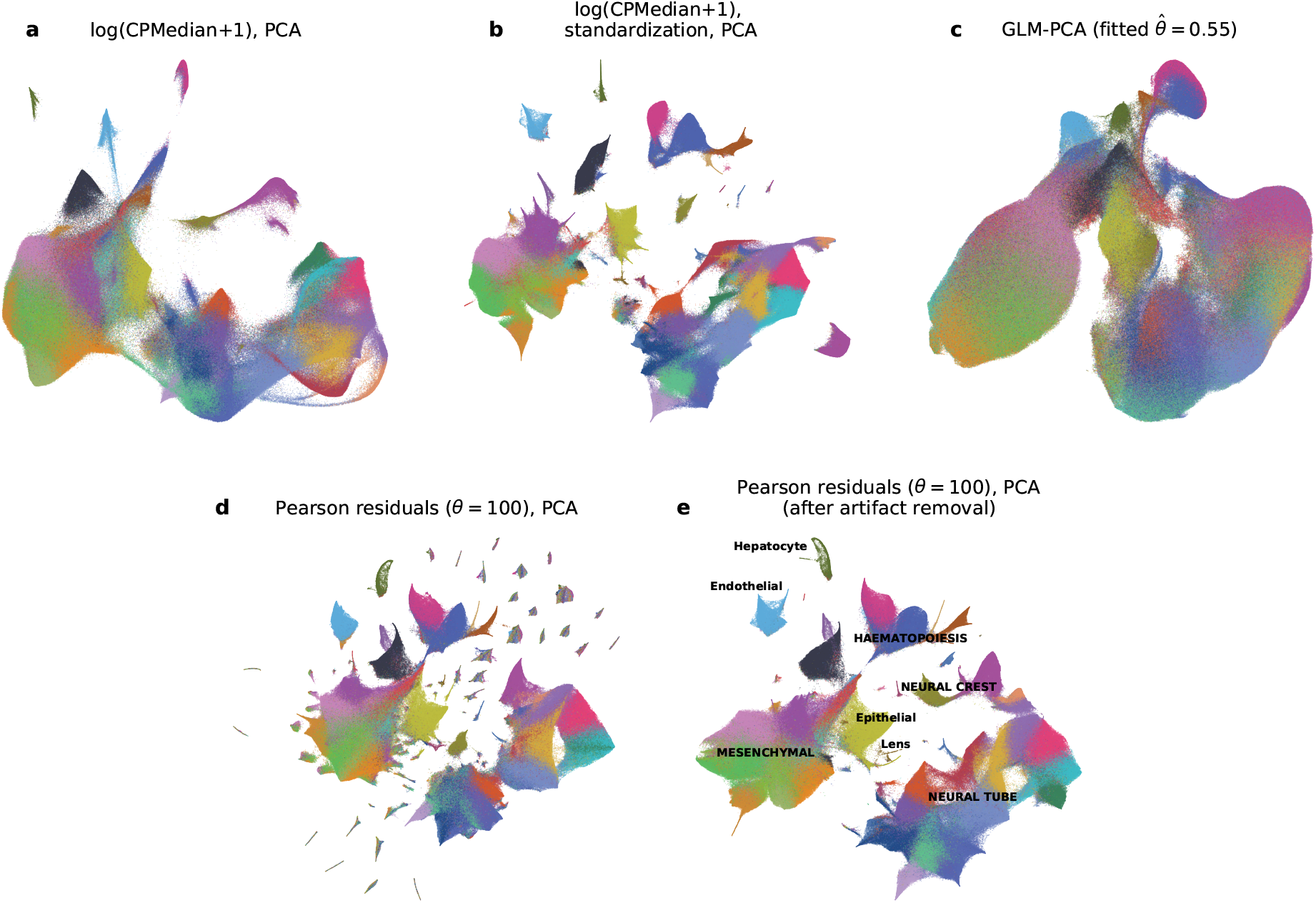
t-SNE embeddings of the organogenesis dataset. All panels show t-SNE embeddings of the organogenesis dataset [39] (2 058 652 cells), colored by the 38 main clusters identified by the original authors. All panels use 2 000 genes with the largest Pearson residual variance. Each panel shows a total of 2 026 641 cells, excluding 32 011 putative doublets identified in the original paper. All t-SNE embeddings were done with exaggeration 4 [38, 41]. **a:** Depth-normalization, median scaling, log-transformation and PCA with 50 principal components. **b:** Same as in (a), but with an additional standardization step that scales the normalized and log-transformed expression of each gene to mean zero and unit variance, as in the original paper [39]. **c:** GLM-PCA with 50 dimensions (NB model with shared overdispersion as a free parameter, estimated to be 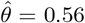). **d:** Analytic Pearson residuals with *θ* = 100 and PCA with 50 principal components. The scattered small islands do not belong to single clusters but instead are spuriously enriched in single embryos. **e:** Same as in (d), but after removing batch-effect genes (Methods). Text labels correspond to the developmental trajectories identified in the original paper [39] (uppercase: multi-cluster trajectories, lowercase: single-cluster trajectories).

However, on closer inspection, embeddings based on Pearson residuals consistently outperformed the other two. For example, while the Pearson residual embeddings clearly separated fine cell types in the full-retina dataset [33], the square-root embedding mixed some of them (we observed the same when using the log-transform). For the same dataset, GLM-PCA embedding did not fully separate some of the biologically distinct cell types. Furthermore, GLM-PCA embeddings often featured Gaussian-shaped blobs with no internal structure (Figure 3), suggesting that some fine manifold structure was lost, possibly due to convergence difficulties.

Embedding the organogenesis dataset [39] using Pearson residuals uncovered a strong and surprising batch artifact: hitherto unnoticed, several genes were highly expressed exclusively in small subsets of cells, with each subset coming from a single embryo. These subsets appeared as isolated islands in the t-SNE embedding (Figure 4), allowing us to uncover and remove this batch effect (Additional File 1: Figure S6), leading to the final, biologically interpretable embedding (Figure 4). In contrast, embeddings based on log-transform or GLM-PCA did not show this batch artifact at all. GLM-PCA took days to converge (Table S1) and could recover only the coarse structure of the data. Interestingly, the final embedding based on Pearson residuals was broadly similar to the embedding obtained after log-transform and standardization of each gene, as expected given that Pearson residuals stabilize the variance by construction (Figure 4). Together, these qualitative observations suggest that analytic Pearson residuals can represent small, distinct subpopulations in large datasets better than other methods.

To quantify the performance of dimensionality reduction methods, we performed a systematic bench-mark using the `Zhengmix8eq` dataset with known ground truth labels [42] (Figure 5). This dataset consists of PBMC cells FACS-sorted into eight different cell types before sequencing [43], with eight types occurring in roughly equal proportions. To make the setup more challenging, we added 10 pseudo-genes expressed only in a group of 50 cells, effectively creating a ninth, rare, cell type (see Methods). We used six methods to select 2000 HVGs (and additionally omitted HVG selection) and ten methods for data transformation and dimensionality reduction to 50 dimensions. We assessed the resulting (6 + 1) · 10 = 70 pipelines using kNN classification of cell types. We used the macro F1 score (harmonic mean between precision and recall, averaged across classes) because this metric fairly averages classifier performance across classes of unequal size. Together, the F1 score of the kNN classifier quantifies how well each pipeline separated cell types in the 50-dimensional representation (Figure 5c). We did not include approaches that use depth normalization with inferred size factors [44] in this comparison.

**Figure 5:**
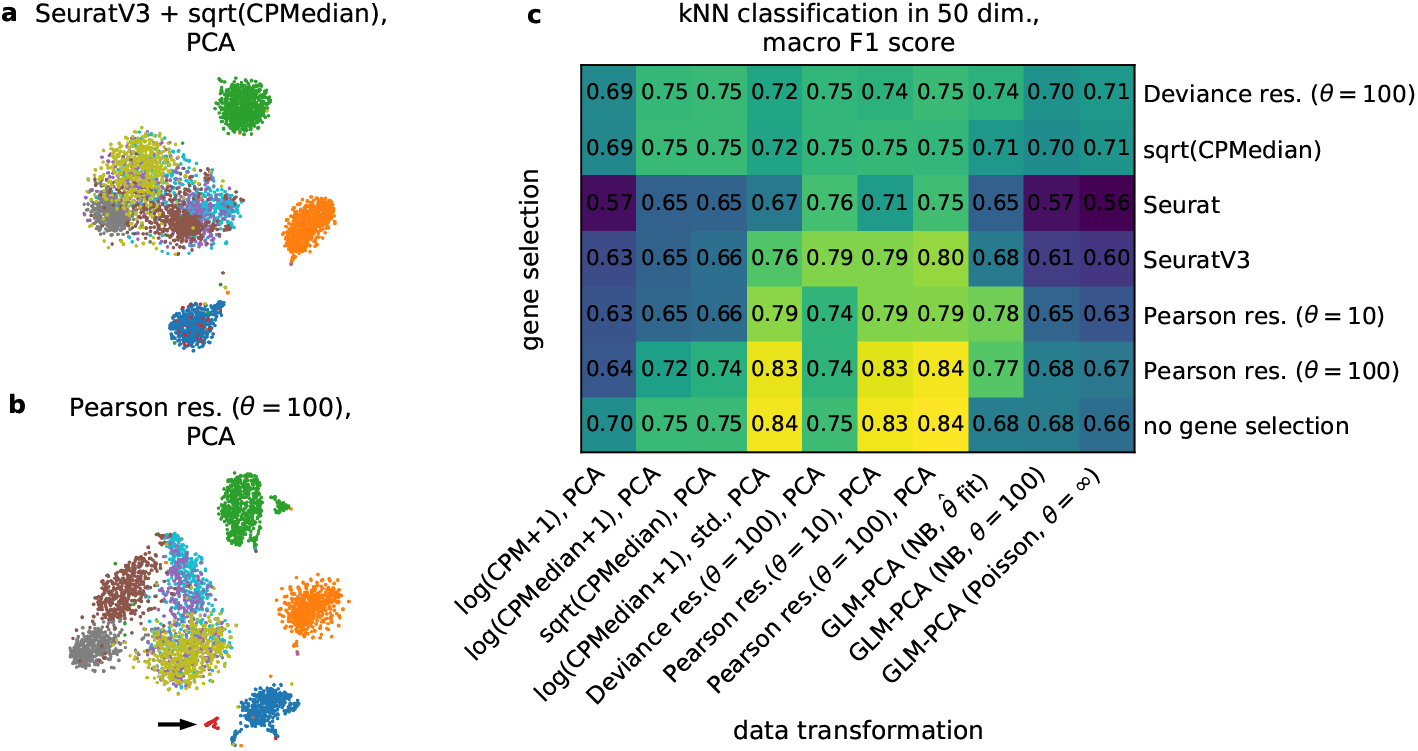
Benchmarking the effect of normalization on cell type separation in reduced dimensionality. We used the Zhengmix8eq dataset with eight ground truth FACS-sorted cell types [42, 43] (3 994 cells) and added ten pseudo-genes expressed in a random group of 50 cells from one type. All HVG selection methods were set up to select 2 000 genes, and all normalization and dimensionality reduction methods reduced the data to 50 dimensions. For details see Methods. **a:** t-SNE embedding after the seurat v3 HVG selection as implemented in Scanpy, followed by depth-normalization, median scaling, square-root transform, and PCA. Colors denote ground truth cell types, the artificially added type is shown in red. **b:** t-SNE embedding after HVG selection by Pearson residuals (*θ* = 100), followed by transformation to Pearson residuals (*θ* = 100), and PCA. Black arrow points at the artificially added type. **c:** Macro F1 score (harmonic mean between precision and recall, averaged across classes to counteract class imbalance) for kNN classification (*k* = 15) of nine ground truth cell types for each of the 70 combinations of HVG selection and data transformation approaches.

The pipeline that used analytic Pearson residuals for both gene selection and data transformation outperformed all other pipelines with respect to cell type classification performance. In contrast, popular methods for HVG selection (e.g. `seurat v3` as implemented in `Scanpy` [5, 35]) combined with log or square-root transformations after depth normalization performed worse and in particular were often unable to separate the rare cell type (Figure 5a,b; see Additional File 1: Figure S7 for additional embeddings). The performance of GLM-PCA was also poor, likely due to convergence issues (with 15-dimensional, and not 50-dimensional, output spaces, GLM-PCA performed on par with Pearson residuals; data not shown), in agreement with what we reported above for the retinal datasets. Finally, deviance residuals [2] were clearly outperformed by Pearson residuals both as gene selection criterion and as data transformation. This is due to the reduced sensitivity of deviance residuals to low-or medium-expression genes (Additional File 1: Figure S2). Note that in terms of the overall classification accuracy no pipeline outperformed Pearson residuals but many pipelines performed similarly well; this is because overall accuracy is not sensitive to the rare cell type, unlike the macro F1 score.

For this dataset, not using gene selection at all performed similarly well to HVG selection using Pearson residuals (Figure 5c), but in general HVG selection is a recommended step in scRNA-seq data analysis [3, 4] and here Pearson residuals performed the best. Also, log-transformed counts that were standardized performed similarly well to Pearson residuals (Figure 5c), in agreement with the above observations on the organogenesis dataset (Figure 4b). Nevertheless, the same organogenesis example showed that Pearson residuals can be more sensitive (Figure 4b, d).

### 2.8 Analytic Pearson residuals are fast to compute

The studied normalization pipelines differ in both space and time complexity. UMI count data are typically very sparse (e.g. in the PBMC dataset, 95% of entries are zero) and can be efficiently stored as a sparse matrix object. Sequencing depth normalization and square-root or log-transformation do not affect the zeros, preserving the sparsity of the matrix, and PCA can be run directly on a sparse matrix. In contrast, Pearson residuals form a dense matrix without any zeros, and so can take a large amount of memory to store (4.5 Gb for the PBMC dataset). For large datasets this can become prohibitive (but note that a smart implementation may be able to avoid storing a dense matrix in memory [45]). In contrast, GLM-PCA can be run directly on a sparse matrix but takes a long time to converge (Table S1), becoming prohibitively slow for bigger datasets.

Computational complexity can be greatly reduced if gene selection is performed in advance. After selecting 1000 genes, Pearson residuals do not require a lot of memory (0.3 Gb for the PBMC dataset) and so can be conveniently used. Note that the Pearson residual variance can be computed per gene, without storing the entire residual matrix in memory. GLM-PCA, however, remained slow even after gene selection (4 h vs. 4 s for Pearson residuals for the PBMC dataset; 2 days vs. 4 m for the organogenesis dataset; Table S1).

## 3 Discussion

We reviewed and contrasted different methods for normalization of UMI count data. We showed that without post-hoc smoothing, the negative binomial regression model of Hafemeister & Satija [1] exhibits high variance in its parameter estimates because it is overspecified, which is why it had to be smoothed in the first place. We argued that instead of smoothing an overspecified model, one should resort to a more parsimonious and theoretically motivated model specification involving an offset term. This made the model equivalent to the rank-one GLM-PCA of Townes et al. [2] and yielded a simple analytic solution, closely related to *correspondence analysis* [17]. Further, we showed that the estimates of pergene overdispersion parameter *θ*_*g*_ in the original paper exhibit substantial and systematic bias. We used negative control datasets from different experimental protocols to show that UMI counts have low overdispersion and technical variation is well described by *θ* ≈ 100 shared across genes.

We found that the approach developed by Hafemeister & Satija [1] and implemented in the R package `scTransform` in practice yields Pearson residuals that are often similar to our analytic Pearson residuals with fixed overdispersion parameter (Additional File 1: Figure S2). We argue that our model with its analytic solution is attractive for reasons of parsimony, theoretical simplicity, and computational speed. Moreover, it provides an explanation for the linear trends in the smoothed estimates in the original paper. We have integrated Pearson residuals into upcoming `Scanpy` 1.9 [5].

Following our manuscript, `scTransform` was updated to `scTransform v2` and now uses the offset model formulation [46]. At the same time, the authors argue that the dependence of the overdispersion parameter *θ*_*g*_ on the gene expression strength is not entirely explained by the estimation bias. To reduce the bias, `scTransform v2` uses `glmGamPoi` [47] to estimate the offsets *β*_0*g*_ and the overdispersion parameters *θ*_*g*_ (which are then smoothed). The authors also refer to the bulk RNA-seq literature, where it has been observed that the overdispersion parameter grows monotonically with gene expression [6, 48, 49]. Given the difficulties with estimating overdispersion for low expression means (see above), we believe that this question requires further investigation. However, as argued above, whether *θ* is assumed to be constant or is allowed to vary between genes, has very little effect on the resulting Pearson residuals.

A parallel publication [50] suggested a Bayesian procedure named `Sanity` for estimating expression strength underlying the observed UMI counts, based on Poisson likelihood and Bayesian shrinkage. Importantly, Pearson residuals are not aiming at estimating the underlying expression strength; rather, they quantify how strongly each observed UMI count deviates from the null model of constant expression across cells. These two approaches can have opposite effects on markers genes of rare cell types: the Bayesian procedure shrinks their expression towards zero whereas our approach yields large Pearson residuals. We argued here that this emphasis on rare cell types is useful for many downstream tasks, but if the interest lies in true expression, approaches like `Sanity` may be more appropriate. Future work should perform comprehensive benchmarks on a variety of tasks [51].

On the practical side, we showed that Pearson residuals outperform other methods for selecting biologically variable genes. They are also better than other pre-processing methods for downstream analysis: in a systematic benchmarking effort, we demonstrated that Pearson residuals provide a good basis for general-purpose dimensionality reduction and for constructing 2D embeddings of single-cell UMI data. In particular, they are well suited for identifying rare cell types and their genetic markers. Applying gene selection prior to dimensionality reduction reduces the computational cost of using Pearson residuals down to negligible. We conclude that analytic Pearson residuals provide a theory-based, fast, and convenient method for normalization of UMI datasets.

## 4 Methods

### 4.1 Mathematical details

#### Analytic solution

The log-likelihood for the model defined in Eqs. 1−2

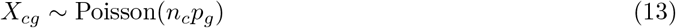

can be, up to a constant, written as

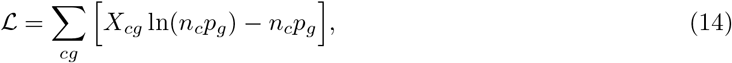

where we used the Poisson density *p*(*x*) = *e*^*x*^*e*^*−μ*^*/x*!. Taking partial derivatives with respect to *n*_*c*_ and *p*_*g*_ and setting them to zero, one obtains

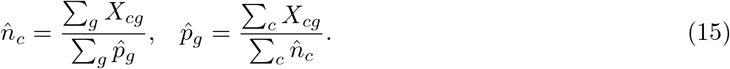

This is a family of solutions. Setting 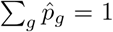, we obtain Eq. 3 and the formulas for 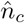 and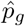 given in Section 2.1.

This derivation does not generalize for the negative binomial model with density

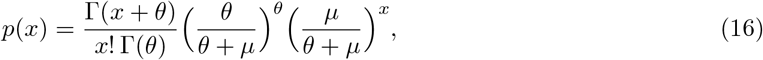

where the log-likelihood (for fixed *θ*), up to a constant, is

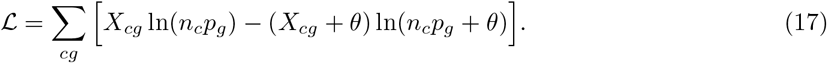

This does not have an analytic maximum likelihood solution. However, for large *θ* values Eq. 3 can be taken as an approximate solution.

#### Deviance residuals

Deviance is defined as the doubled difference between the log-likelihood of the saturated model and the log-likelihood of the actual model. The saturated model, in our case, is a full rank model with 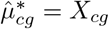.

For the Poisson model, the deviance can therefore be obtained from Eq. 14 and is equal to

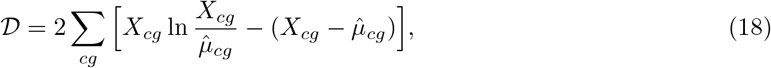

where the terms with 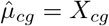 are taken to be zero.

Deviance residuals are defined as square roots of the respective deviance terms, such that the sum of squared deviance residuals is equal to the deviance (note that for the Gaussian case this already holds true for the raw residuals, because the saturated model has zero log-likelihood, and the deviance is simply the squared error). It follows that for the Poisson model deviance residuals [2] are given by

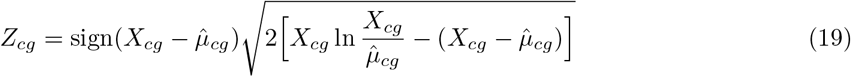

Similarly, for the negative binomial model with fixed *θ*, the deviance residuals follow from Eq. 17 and are given by

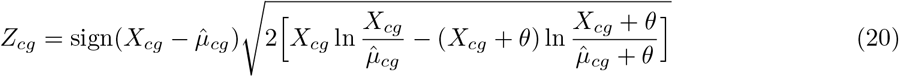

It is easy to verify that this formula reduces to the Poisson case when *θ → ∞*. When computing deviance residuals, we estimated 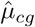 using Eq. 3.

#### Clipping Pearson residuals

Clipping Pearson residuals to 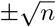 as suggested by Hafemeister & Satija [1] is needed to avoid large residual variance in rarely expressed genes (Additional File 1: Figure S2d). The intuition behind this heuristic is as follows. Consider a UMI dataset with *n* cells containing a biologically distinct rare population *P* of size *m ≪ n*. Let this population have a marker gene with expression following Poisson(*λ*) for the cells from *P*, and zero expression for all *n* − *m* remaining cells. For simplicity we assume the Poisson model here, and further assume that all cells have the same sequencing depth.

The expected average expression of this gene is *λm/n* and so the expected Pearson residual value for this gene for the cells from *P* is 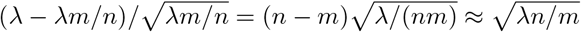.

With the clipping threshold 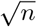, clipping will happen whenever *λ > m*, i.e. when the population P is either very small or has very large UMI counts. For example, a population of 10 cells having a marker gene with the within-population mean expression of 20 UMIs, will result in clipped residuals, as if the within-population mean expression were ∼10 UMIs. This may have a large effect on the leading principal components (even PC1) if the data contain a very small number of cells with strong marker gene expression.

#### Pearson residuals of biologically variable genes

It is instructive to observe the effect Pearson residuals have on genes that have the same variance of log-expression but different expression means. Consider a gene that has expression *μ* in half of the cells and is upregulated by a factor of two in the other half of the cells. Then its expression mean is 1.5*μ*, and the Pearson residuals are close to 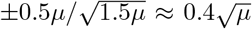, i.e. the variance of Pearson residuals grows linearly with *μ*. This makes sense because for higher-expressed genes there is more statistical certainty about over-Poisson variability, but at the same time highlights that Pearson residuals do not aim to estimate the underlying (log-)expression, unlike e.g. `Sanity` [50].

### 4.2 Experimental details

#### Analyzed datasets and preprocessing

Used datasets are listed in Table S2. For the organogenesis dataset and the FACS-sorted PBMC dataset, we applied no further filtering. In all remaining datasets we excluded genes that were expressed in fewer than 5 cells, following Hafemeister & Satija [1]. The data were downloaded following links in original publications in form of UMI count tables. Direct links to all data sources are given in our Github repository https://github.com/berenslab/umi-normalization.

#### HVG selection

For gene selection using `sqrt(CPMedian)`, Pearson residuals, and deviance residuals, we applied the respective data transformation and used the variance after transformation as selection criterion. For `Seurat` and `Seurat v3` methods, we used the respective `Scanpy` implementations. In brief, these two methods regress out the mean-variance relationship, and return an estimate of the ‘excess’ variance for each gene [34, 35]. For `scTransform` we used the corresponding R package [1]. The Fano factor was computed after normalizing by sequencing depth and scaling by median sequencing depth.

#### Data transformation and dimensionality reduction

We used the following abbreviations to denote data transformations: `sqrt(CPMedian)` — normalization by sequencing depth, followed by scaling by the median depth across all cells (‘counts per median’), followed by the square-root transform; `log(CPMedian + 1)` — normalization by sequencing depth, followed by scaling by the median depth across all cells, followed by log(*x* + 1) transform; `log(CPMedian + 1) + standardization` — same as `log(CPMedian + 1)`, but followed by centering each gene at mean zero and unit variance; `log(CPM + 1)` — normalization by sequencing depth, followed by scaling by one million (‘counts per million’), followed by log(*x* + 1) transform. Pearson residuals were computed with Eq. 12 and then clipped to 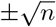. Deviance residuals were computed with Eq. 20.

All of these methods were typically followed by dimensionality reduction by PCA to 50 dimensions using the `Scanpy` implementation [5], unless otherwise stated.

Further, we used three variants of GLM-PCA to transform raw counts and reduce dimensionality down to 50 in a joint step: Poisson GLM-PCA, negative binomial GLM-PCA with estimation of single overdispersion parameter *θ* shared across genes, and negative binomial GLM-PCA with fixed shared *θ*. In Townes et al. [2], the authors only used the former two methods. Whenever possible, we used the `glmpca-py` implementation with default settings. When we reduced the PBMC dataset to 1 000 genes for Additional File 1: Figure S4f, GLM-PCA did not converge with default penalty 1, so we increased it to 5, following the tuning procedure used in the authors’ `R` implementation. Similarly, negative binomial GLM-PCA with estimation of *θ* did not converge on the benchmark dataset (Figure 5) when we used gene selection by either Deviance residuals (*θ* = 100) or Pearson residuals (*θ* = 10). For these two cases, we had to increase the penalty to 10. On the organogenesis dataset, the Python implementation did not converge within reasonable time, so for this dataset, we resorted to the R implementation. It uses a different optimization method and employs stochastic minibatches. All reported GLM-PCA results for this dataset are for batchsize 10 000 as batchsizes 100 and 1 000 (default) resulted in considerably longer runtimes. Because the R implementation does not support NB GLM-PCA with fixed theta, for this dataset we used GLM-PCA with jointly fit 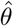.

Unless otherwise stated, all residuals and GLM-PCA with fixed *θ* used *θ* = 100. Whenever gene selection was performed prior to a data transformation that required sequencing depths, we computed those depths using the sum over selected genes only.

#### Benchmarking cell type separation with kNN classification

We used the `Zhengmix8eq` dataset with known ground truth labels obtained by FACS-sorting of eight PBMC cell types [42, 43]. There were 400−600 cells in each cell type. We created a ninth, artificial population from 50 randomly selected B-cells (marked blue in Figure 5). To mimic a separate cell type, we added 10 pseudo marker that had zero expression everywhere apart from those 50 cells. For those 50 cells, UMI values were simulated as Poisson(*n*_*i*_*p*), where *n*_*i*_ is the sequencing depth of the *i*-th selected cell (range: 452−9697), and expression fraction *p* was set to 0.001.

We then applied the 70 normalization pipelines shown in Figure 5 to this dataset. Each pipeline either used one of the six methods to select 2000 HVGs or proceeded without HVG selection, followed by one of the ten methods for data transformation and dimensionality reduction to 50 dimensions. To assess cell type separation in this output space, we used a kNN classifier with a leave-one-out cross-validation procedure: For each cell, we trained a kNN classifier on the remaining *n* − 1 cells. This resulted in a class prediction for each cell based on the majority vote of its *k* = 15 neighboring cells. We quantified the performance of this prediction by computing the macro F1 score (harmonic mean between precision and recall, averaged across classes to counteract class imbalance). We used the `sklearn` implementations for kNN classification and the F1 score [52].

#### Measuring runtimes

All runtimes given in Table S1 are wall times from running the code in a Docker container with an Ubuntu 18 system on a machine with 256 GB RAM and 2 × 24 CPUs at 2.1 Ghz (Xeon Silver 4116 Dodecacore). The Docker container was restricted to use at most 30 CPU threads. To reduce overhead, we did not use `Scanpy` for timing experiments, and instead used `numpy` for basic computations and `sklearn` for PCA with default settings. Note that we used different implementations of GLM-PCA for the PBMC and organogenesis dataset (see above for details).

#### t-SNE embeddings

All t-SNE embeddings were made following recommendations from a recent paper [38] using the FIt-SNE implementation [53]. We used the PCA (or, when applicable, GLM-PCA) representation of the data as input. We used default FIt-SNE parameters, including automatically chosen learning rate. For initialization, we used the first two principal components of the data, scaled such that PC1 had standard deviation 0.0001 (as is default in FIt-SNE). The initialization was shared among all embeddings shown in the same figure, i.e. PCs of one data representation were used to initialize all other embeddings as well. For all datasets apart from the organogenesis one, we used perplexity combination of 30 and *n/*100, where *n* is the sample size [38]. For the organogenesis dataset embeddings we used perplexity 30 and exaggeration 4 [41].

## Availability of Data and Materials

All software needed to reproduce the analysis and figures presented in this work are published under GNU Affero General Public License v3.0 on Github at https://www.github.com/berenslab/umi-normalization [54]. The state of the repository at submission of this manuscript is archived at https://zenodo.org/record/5150534 [55].

All datasets used in this work are publicly available and listed in the following Table S2. Detailed download instructions can be found at our Github repository.

## Acknowledgements

We thank Christoph Hafemeister, Rahul Satija, William Townes, Florian Wagner, Constantin Ahlmann-Eltze, and Erik van Nimwegen for discussions and helpful comments, and the `Scanpy` team for support.

## Ethics approval

Not applicable.

## Supplementary information

Additional File 1 contains supplementary Figures S1−S7

---

**Table S1:** Runtimes for different normalization pipelines. The datasets are: the full 33k PBMC dataset, the PBMC dataset after selecting 1 000 HVGs, and the organogenesis dataset [39] after selecting 2 000 HVGs. Genes with largest Pearson residual variances were selected, which took 9 seconds (PBMC) and 15 minutes (organogenesis), respectively. See Methods for details. All runtimes measured on a machine with 256 Gb RAM and 30 CPU threads at 2.1 GHz.

|  | Sqrt(CPMedian), PCA | Pearson residuals, PCA | GLM-PCA |
| --- | --- | --- | --- |
| 33k PBMC | 31 s | 35 s | 15 h |
| 33k PBMC, 1000 HVGs | 3 s | 4 s | 4 h |
| 2M organogenesis, 2000 HVGs | 166 s | 224 s | 52 h |

**Table S2:** Overview of UMI datasets used for analysis. In the 10X control dataset [56], we used only sample 1. In the MicrowellSeq control dataset [58], we used the E14 dataset. In the three retinal datasets [33, 36, 37], we only used cells from the largest batch. The FACS-sorted PBMC dataset was assembled by authors of a recent paper[42], based on a benchmarking dataset published earlier [43]. Numbers of genes and cells are after batch selection (where applicable) and initial gene filtering (see Methods). Scripts performing these operations and detailed download instructions for all materials are published in our Github repository at http://www.github.com/berenslab/umi-normalization. The accession numbers refer to archived datasets at the Gene Expression Omnibus (NCBI), the Sequence Read Archive (NCBI), or ArrayExpress (EMBL-EBI). (*): Data directly obtained from 10X Genomics at https://support.10xgenomics.com/single-cell-gene-expression/datasets/1.1.0/pbmc33k. (**): The accession number links to the base dataset that the original authors used to construct the ground-truth dataset for their paper [42]. To obtain the dataset used here, use the Bioconductor 3.1.3 R package DuoClustering2018 or visit the authors’ website (http://imlspenticton.uzh.ch/robinson_lab/DuoClustering2018/).

| reference | accession no. | protocol | cells | genes | species | description |
| --- | --- | --- | --- | --- | --- | --- |
| 33k PBMC | * | 10X v1 | 33 148 | 16 809 | human | peripheral blood mononuclear cells |
| [43] | SRP073767** | 10X v1 | 3 994 | 15 715 | human | FACS-sorted PBMC cells |
| [56] | E-MTAB-5480 | 10X v2 | 2 000 | 13 025 | human | droplets of bulk RNA solution |
| [57] | GSM1599501 | inDrop | 953 | 25 025 | human | droplets of bulk RNA solution |
| [58] | GSM2906413 | MicrowellSeq | 9 994 | 15 069 | mouse | non-differentiating stem cells |
| [33] | GSE63472 | DropSeq | 24 769 | 17 973 | mouse | retinal cells |
| [36] | GSE81904 | DropSeq | 13 987 | 16 520 | mouse | retinal bipolar cells |
| [37] | GSE133382 | 10X v2 | 15 750 | 17 685 | mouse | retinal ganglion cells |
| [39] | GSE119945 | sci-RNA-seq3 | 2 058 652 | 26 183 | mouse | organogenesis of mouse embryo cells |

## References

[1] Hafemeister C, Satija R. Normalization and variance stabilization of single-cell RNA-seq data using regularized negative binomial regression. Genome Biology. 2019;20:296.

[2] Townes FW, Hicks SC, Aryee MJ, Irizarry RA. Feature selection and dimension reduction for single-cell RNA-Seq based on a multinomial model. Genome Biology. 2019;20:295.

[3] Luecken MD, Theis FJ. Current best practices in single-cell RNA-seq analysis: a tutorial. Molecular Systems Biology. 2019;15(6).

[4] Amezquita RA, Lun AT, Becht E, Carey VJ, Carpp LN, Geistlinger L, et al. Orchestrating single-cell analysis with Bioconductor. Nature Methods. 2020;17(2):137–145.

[5] Wolf FA, Angerer P, Theis FJ. SCANPY: large-scale single-cell gene expression data analysis. Genome Biology. 2018;19(1):1–5.

[6] Love MI, Huber W, Anders S. Moderated estimation of fold change and dispersion for RNA-seq data with DESeq2. Genome Biology. 2014;15(12):1–21.

[7] Eling N, Richard AC, Richardson S, Marioni JC, Vallejos CA. Correcting the mean-variance dependency for differential variability testing using single-cell RNA sequencing data. Cell Systems. 2018;7(3):284–294.

[8] Lopez R, Regier J, Cole MB, Jordan MI, Yosef N. Deep generative modeling for single-cell transcriptomics. Nature Methods. 2018;15(12):1053–1058.

[9] Svensson V, Gayoso A, Yosef N, Pachter L. Interpretable factor models of single-cell RNA-seq via variational autoencoders. Bioinformatics. 2020;.

[10] Sarkar A, Stephens M. Separating measurement and expression models clarifies confusion in singlecell RNA sequencing analysis. Nature Genetics. 2021;53(6):770–777.

[11] Grün D, Kester L, Van Oudenaarden A. Validation of noise models for single-cell transcriptomics. Nature methods. 2014;11(6):637–640.

[12] Svensson V. Droplet scRNA-seq is not zero-inflated. Nature Biotechnology. 2020;38(2):147–150.

[13] Agresti A. Foundations of linear and generalized linear models. John Wiley & Sons; 2015.

[14] Culhane A. Correspondence Analysis in R. GitHub; 2020. https://aedin.github.io/PCAworkshop/articles/c_COA.html.

[15] Hill MO. Correspondence analysis: a neglected multivariate method. Journal of the Royal Statistical Society: Series C (Applied Statistics). 1974;23(3):340–354.

[16] Greenacre M, Hastie T. The geometric interpretation of correspondence analysis. Journal of the American Statistical Association. 1987;82(398):437–447.

[17] Greenacre M. Correspondence analysis in practice. Chapman and Hall/CRC; 2007.

[18] Holmes S. Multivariate data analysis: the French way. In: Probability and statistics: Essays in honor of David A. Freedman. Institute of Mathematical Statistics; 2008. p. 219–233.

[19] Hirschfeld HO. A connection between correlation and contingency. In: Mathematical Proceedings of the Cambridge Philosophical Society. vol. 31. Cambridge University Press; 1935. p. 520–524.

[20] Willson L, Folks J, Young J. Complete sufficiency and maximum likelihood estimation for the two-parameter negative binomial distribution. Metrika. 1986;33(1):349–362.

[21] Clark SJ, Perry JN. Estimation of the negative binomial parameter κ by maximum quasi-likelihood. Biometrics. 1989;p. 309–316.

[22] Lord D. Modeling motor vehicle crashes using Poisson-gamma models: Examining the effects of low sample mean values and small sample size on the estimation of the fixed dispersion parameter. Accident Analysis & Prevention. 2006;38(4):751–766.

[23] Lord D, Miranda-Moreno LF. Effects of low sample mean values and small sample size on the estimation of the fixed dispersion parameter of Poisson-gamma models for modeling motor vehicle crashes: A Bayesian perspective. Safety Science. 2008;46(5):751–770.

[24] Kim JK, Kolodziejczyk AA, Ilicic T, Teichmann SA, Marioni JC. Characterizing noise structure in single-cell RNA-seq distinguishes genuine from technical stochastic allelic expression. Nature Communications. 2015;6(1):1–9.

[25] Wang J, Huang M, Torre E, Dueck H, Shaffer S, Murray J, et al. Gene expression distribution deconvolution in single-cell RNA sequencing. Proceedings of the National Academy of Sciences. 2018;115(28):E6437–E6446.

[26] Lopez-Delisle L, Delisle JB. baredSC: Bayesian Approach to Retrieve Expression Distribution of Single-Cell. bioRxiv. 2021;.

[27] Bar-Lev SK, Enis P. On the classical choice of variance stabilizing transformations and an application for a Poisson variate. Biometrika. 1988;75(4):803–804.

[28] Anscombe FJ. The transformation of Poisson, binomial and negative-binomial data. Biometrika. 1948;35(3/4):246–254.

[29] Freeman MF, Tukey JW. Transformations related to the angular and the square root. The Annals of Mathematical Statistics. 1950;p. 607–611.

[30] Wagner F. Straightforward clustering of single-cell RNA-Seq data with t-SNE and DBSCAN. BioRxiv. 2019;p. 7703.8.

[31] Wagner F. Monet: An open-source Python package for analyzing and integrating scRNA-Seq data using PCA-based latent spaces. bioRxiv. 2020;.

[32] Warton DI. Why you cannot transform your way out of trouble for small counts. Biometrics. 2018;74(1):362–368.

[33] Macosko EZ, Basu A, Satija R, Nemesh J, Shekhar K, Goldman M, et al. Highly parallel genome-wide expression profiling of individual cells using nanoliter droplets. Cell. 2015;161(5):1202–1214.

[34] Satija R, Farrell JA, Gennert D, Schier AF, Regev A. Spatial reconstruction of single-cell gene expression data. Nature Biotechnology. 2015;33(5):495–502.

[35] Stuart T, Butler A, Hoffman P, Hafemeister C, Papalexi E, Mauck III WM, et al. Comprehensive integration of single-cell data. Cell. 2019;177(7):1888–1902.

[36] Shekhar K, Lapan SW, Whitney IE, Tran NM, Macosko EZ, Kowalczyk M, et al. Comprehensive classification of retinal bipolar neurons by single-cell transcriptomics. Cell. 2016;166(5):1308–1323.

[37] Tran NM, Shekhar K, Whitney IE, Jacobi A, Benhar I, Hong G, et al. Single-cell profiles of retinal ganglion cells differing in resilience to injury reveal neuroprotective genes. Neuron. 2019;104(6):1039– 1055.

[38] Kobak D, Berens P. The art of using t-SNE for single-cell transcriptomics. Nature Communications. 2019;10(1):1–14.

[39] Cao J, Spielmann M, Qiu X, Huang X, Ibrahim DM, Hill AJ, et al. The single-cell transcriptional landscape of mammalian organogenesis. Nature. 2019;566(7745):496–502.

[40] Lun A. Overcoming systematic errors caused by log-transformation of normalized single-cell RNA sequencing data. BioRxiv. 2018;p. 404962.

[41] Böhm JN, Berens P, Kobak D. A Unifying Perspective on Neighbor Embeddings along the Attraction-Repulsion Spectrum. arXiv preprint 200708902. 2020;.

[42] Duú A, Robinson MD, Soneson C. A systematic performance evaluation of clustering methods for single-cell RNA-seq data. F1000Research. 2018;7.

[43] Zheng GX, Terry JM, Belgrader P, Ryvkin P, Bent ZW, Wilson R, et al. Massively parallel digital transcriptional profiling of single cells. Nature Communications. 2017;8(1):1–12.

[44] Lun AT, Bach K, Marioni JC. Pooling across cells to normalize single-cell RNA sequencing data with many zero counts. Genome Biology. 2016;17(1):1–14.

[45] Irizarry R. R package with Methods for Small Counts Stored in a Sparse Matrix. GitHub; 2021. https://github.com/rafalab/smallcount.

[46] Choudhary S, Satija R. Comparison and evaluation of statistical error models for scRNA-seq. bioRxiv. 2021;.

[47] Ahlmann-Eltze C, Huber W. glmGamPoi: fitting Gamma-Poisson generalized linear models on single cell count data. Bioinformatics. 2020;36(24):5701–5702.

[48] Anders S, Huber W. Differential expression analysis for sequence count data. Genome Biology. 2010;11(10):1–12.

[49] Law CW, Chen Y, Shi W, Smyth GK. voom: Precision weights unlock linear model analysis tools for RNA-seq read counts. Genome Biology. 2014;15(2):1–17.

[50] Breda J, Zavolan M, van Nimwegen E. Bayesian inference of gene expression states from single-cell RNA-seq data. Nature Biotechnology. 2021;p. 1–9.

[51] Ahlmann-Eltze C, Huber W. Transformation and Preprocessing of Single-Cell RNA-Seq Data. bioRxiv. 2021;.

[52] Pedregosa F, Varoquaux G, Gramfort A, Michel V, Thirion B, Grisel O, et al. Scikit-learn: Machine learning in Python. Journal of Machine Learning Research. 2011;12:2825–2830.

[53] Linderman GC, Rachh M, Hoskins JG, Steinerberger S, Kluger Y. Fast interpolation-based t-SNE for improved visualization of single-cell RNA-seq data. Nature methods. 2019;16(3):243–245.

[54] Lause J. Analytic Pearson residuals for normalization of single-cell RNA-seq UMI data. GitHub; 2021. https://github.com/berenslab/umi-normalization.

[55] Lause J. Analytic Pearson residuals for normalization of single-cell RNA-seq UMI data. Zenodo; 2021. https://doi.org/10.5281/zenodo.5150534.

[56] Svensson V, Natarajan KN, Ly LH, Miragaia RJ, Labalette C, Macaulay IC, et al. Power analysis of single-cell RNA-sequencing experiments. Nature Methods. 2017;14(4):381–387.

[57] Klein AM, Mazutis L, Akartuna I, Tallapragada N, Veres A, Li V, et al. Droplet barcoding for single-cell transcriptomics applied to embryonic stem cells. Cell. 2015;161(5):1187–1201.

[58] Han X, Wang R, Zhou Y, Fei L, Sun H, Lai S, et al. Mapping the mouse cell atlas by microwell-seq. Cell. 2018;172(5):1091–1107.

