## Additional File 1, Supplementary figures 1-7 for "Analytic Pearson residuals for normalization of single-cell RNA-seq UMI data"

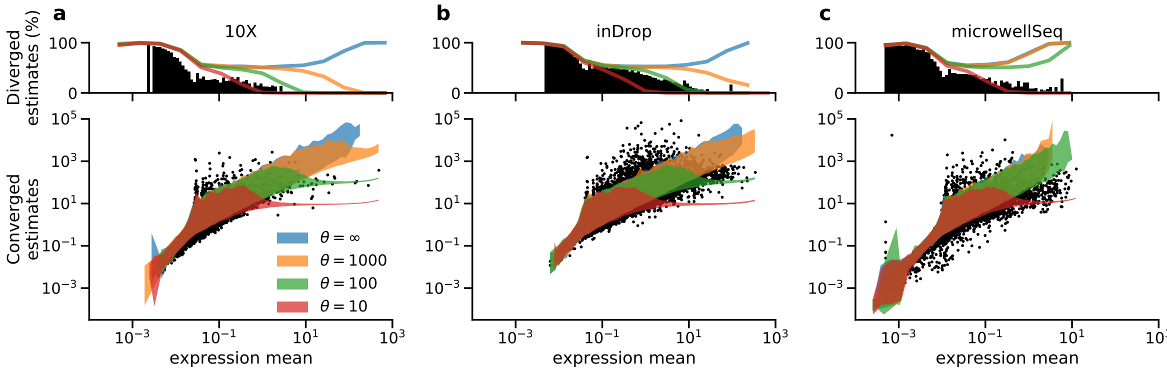


**Figure S1: Overdispersion in negative control datasets.** In the lower panels, each dot corresponds to one gene $g$. The overdispersion parameter estimates were obtained with `theta.ml()` using 100 iterations, using the expression means predicted by our model. Diverged estimates were clipped as in Figure 1g. Colors show the 5th to 95th percentile regions of the overdispersion estimates for simulated datasets with a known $\theta$ shared across genes. For each value of $\theta \in \{10, 100, 1000, \infty\}$, we simulated 1000 datasets according to Eq. 5, using means $\mu_{cg} = p_g n_c$ with empirical $n_c$ and a series of $p_g$ values chosen to cover the entire data range. The upper panels show the fraction of genes for which the estimate diverged (black: original data, colors: simulated data). **a:** 10x Genomics Chromium technical negative control dataset (sample 1), consisting of RNA solution split into droplets [1]. 2 000 cells. **b:** inDrop technical negative control dataset prepared as in (a) [2]. 953 cells. **c:** microwellSeq biological negative control dataset from non-differentiating embryonic stem cell culture (E14 line from strain 129P2/Ola) [3]. 9 994 cells. Biological negative control datasets have been shown to exhibit larger variability (i.e. lower $\theta$) compared to technical negative control datasets [4].

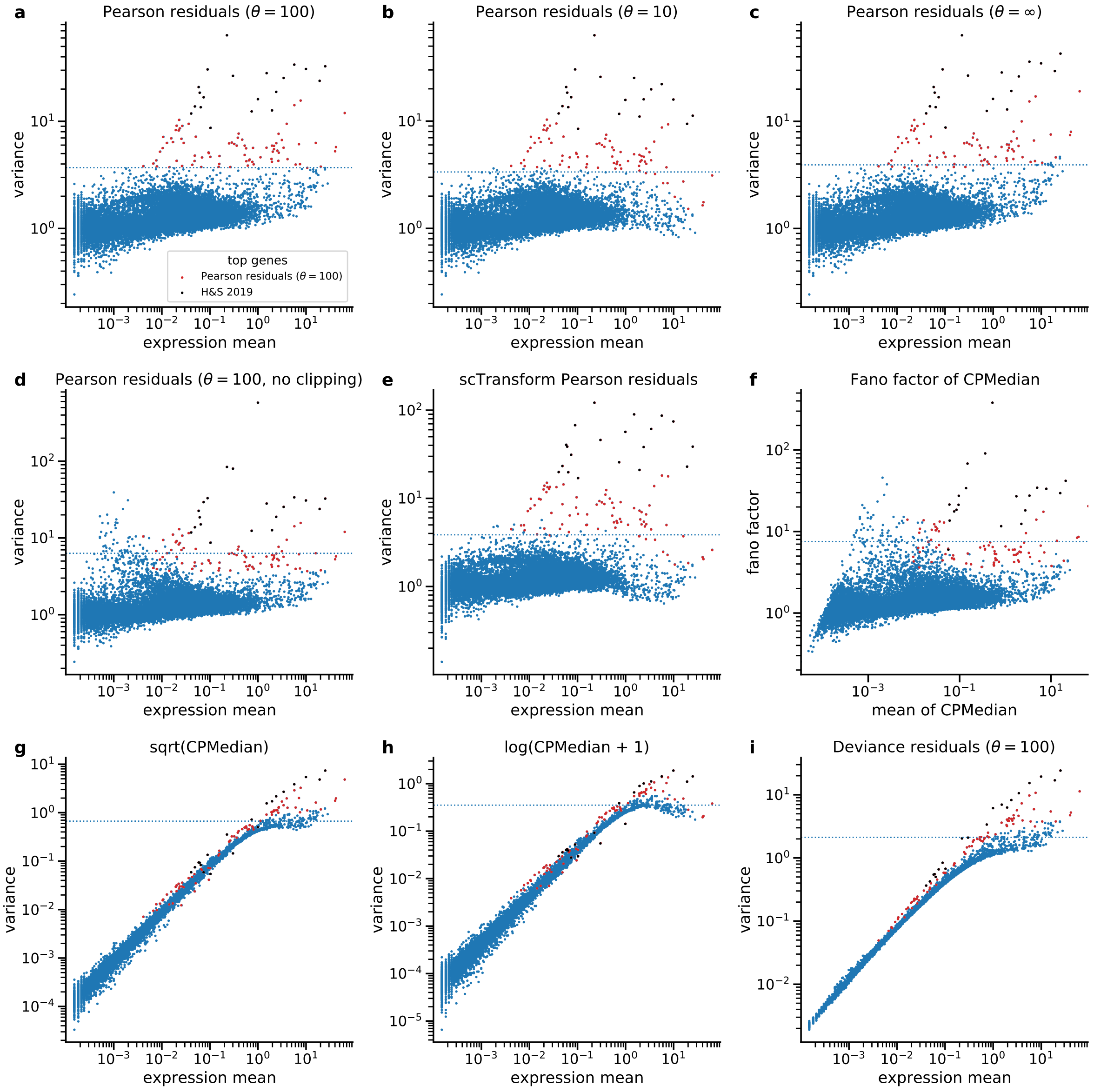


**Figure S2: Comparing gene selection criteria.** Each dot shows the selection criterion for a single gene in the PBMC dataset after applying a normalization method. The 100 genes with the largest criterion values in each panel lie above the dotted horizontal line. Red dots show the top 100 genes selected by the variance of Pearson residuals with $\theta = 100$. Black dots show the top 20 genes selected by `scTransform` [5]. **a:** Reproduced from Figure 2b. **b:** Analytic Pearson residuals with $\theta = 10$, roughly corresponding to `scTransform`. **c:** Analytic Pearson residuals with $\theta = \infty$, i.e. a Poisson model. **d:** Analytic Pearson residuals with $\theta = 100$ but without clipping of the residuals to $\pm\sqrt{n}$. This leads to selection of some low-expression genes. **e:** Pearson residuals of the Hafemeister & Satija [5] model, obtained via the R implementation of `scTransform`. 87% of the top-100, and 83% of the top-1000 genes are the same as in the selection by analytic Pearson residuals with $\theta = 100$. **f:** Fano factor after sequencing depth normalization and median scaling. **g:** Variance after sequencing depth normalization, median-scaling and square-root transform. **h:** Variance after sequencing depth normalization, median-scaling and $\log(1+x)$-transform. Note that here the variance is a non-monotonic function of the average expression. Note also that some biologically variable genes (black and red dots) in both (g) and (h) lie below the main cloud, i.e. have *lower* variance than the genes with the same average expression but without biological variability. This happens for genes that are strongly expressed but only in a small subset of cells. Square-root and log-transformations act 'too strong' on these large counts, yielding lower overall variance. **i:** Variance of deviance residuals for $\theta = 100$. Consistent with Supplementary Figure S7 in Townes et al. [6], only high expression genes are selected.

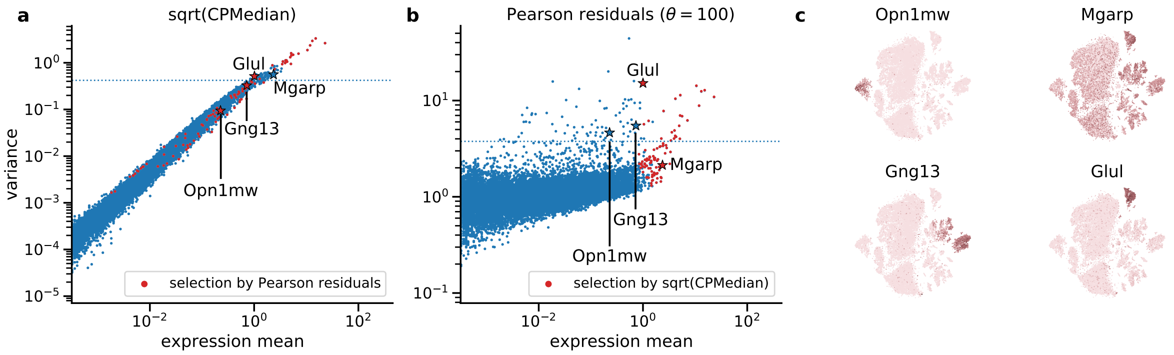


**Figure S3: Selection of variable genes for a retinal dataset.** All panels as in Figure 2, here showing gene selection for the largest batch of the retina dataset from [7]. It consists of replicates `p1` and `r4`–`r6`, with a total of $N = 24\,769$ cells. The cone photoreceptor marker *Opn1mv* and the bipolar cell marker *Gng13* were only selected when the variance of Pearson residuals was used as criterion. The diffusely expressed gene *Mgarp* is present in many cell types, and was only selected based on the variance of the square-root transformed data. The Müller glia marker *Glul* was selected by both methods.

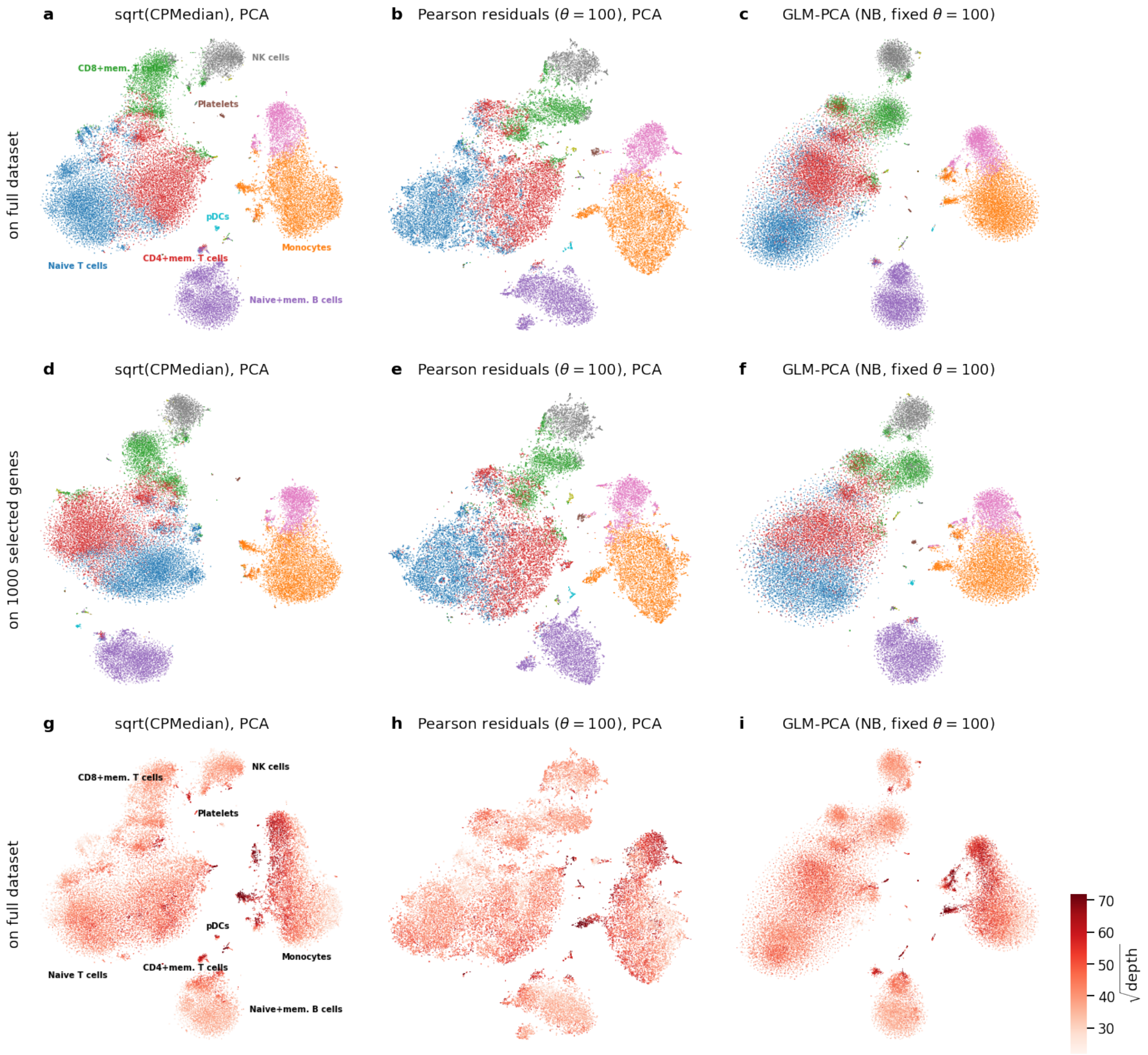


**Figure S4: t-SNE embeddings based on different data transformation approaches.** Each panel shows a t-SNE of the PBMC dataset, the different panels in each row show embeddings based on a different data transformation method with reduction to 50 dimensions (see Methods). Colors correspond to 10 k-means clusters provided by 10x Genomics together with the PBMC dataset. We annotated eight clusters based on known marker genes (taken from [8]). **a–c:** No gene selection was used for these panels. **d–f:** Same as a–c, but using only the 1000 genes with the highest Pearson residual variance. **g–i:** Same as a–c, but overlayed with square-root-transformed sequencing depth. Values above 70 were clipped.

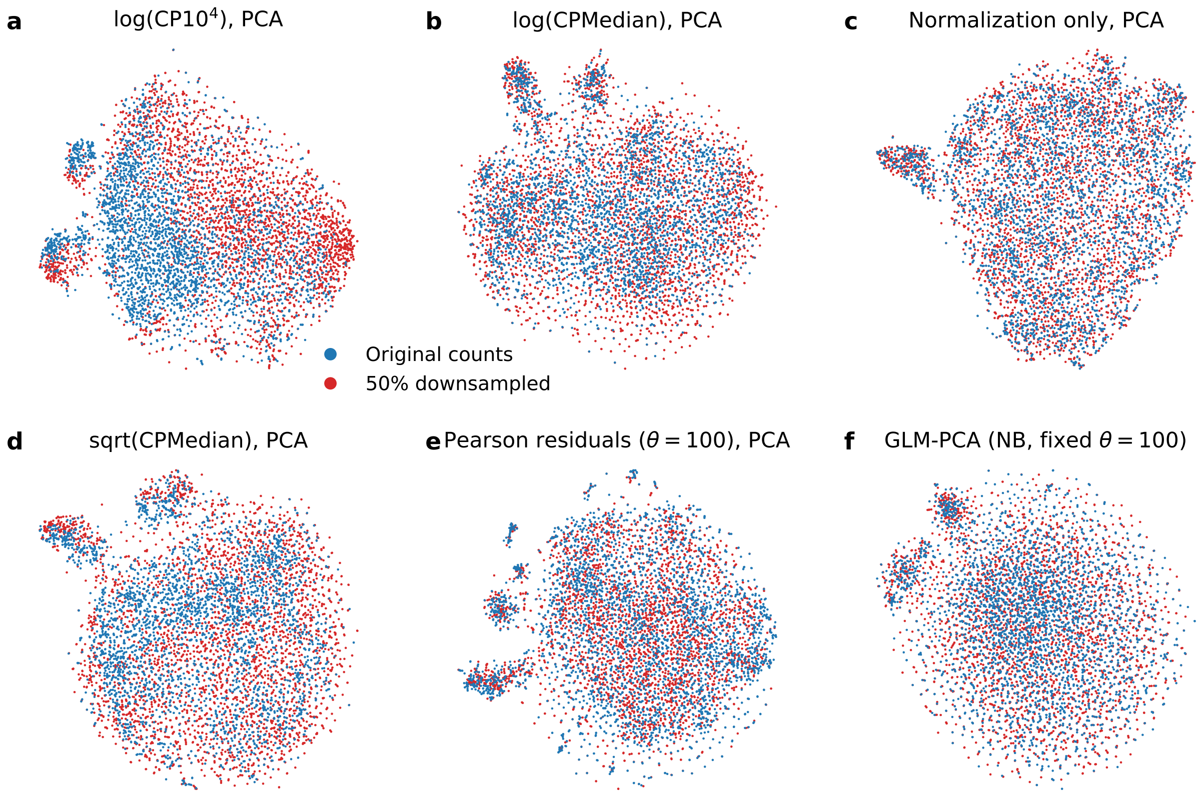


**Figure S5: t-SNE embeddings of the monocyte cluster with downsampling.** We analyzed the monocyte cluster from the PBMC dataset (marked orange in Figure S4, $n = 6\,307$ cells). Half of the cells (randomly selected) were downsampled to 50% of their original sequencing depth, by simulating new counts $\tilde{X}_{cg} = \text{Binomial}(n = X_{cg}, p = 0.5)$, following the analysis in Hafemeister & Satija [5] (their Figure 6C). We then compared how well different normalization strategies remove the simulated batch-effect. PCA and t-SNE were applied after normalization as in Figure S4. **a:** Depth-normalized counts, scaled with $10^4$ and then log-transformed, as in Hafemeister & Satija [5]. Note the strong batch effect due to the large scale factor. **b:** Depth-normalized counts, scaled with median depth ($\approx$ 1500) and then log-transformed. Note that this smaller scale factor was more appropriate and removed the batch effect shown in (a). **c:** Depth-normalized counts. At the cost of not applying a variance-stabilizing transform, no batch effect remains. **d:** Depth-normalized counts, square-root transformed. A weak batch effect remains. **e:** Pearson residuals, computed with $\theta = 100$. **f:** Negative binomial GLM PCA, computed with fixed $\theta = 100$.

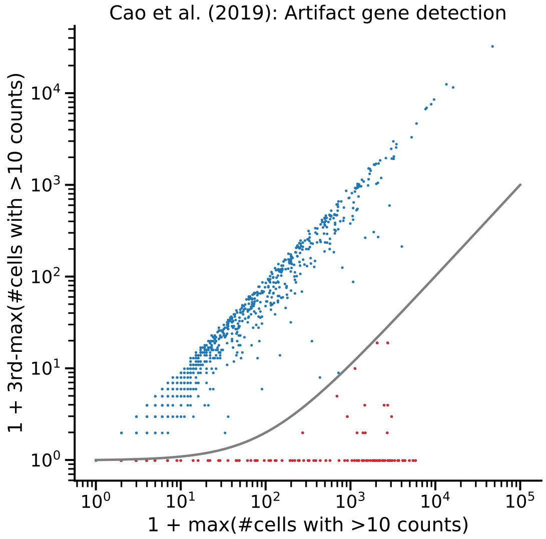


**Figure S6: Artifact genes in the organogenesis dataset.** Scatter plot over the 2 000 highly variable genes that we selected using Pearson residuals. For each gene and each of the 61 embryos in the data, we computed how many cells in that embryo have over 10 UMI counts of that gene. The plot compares the largest number across embryos with the third largest number. For the majority of the genes, these values were very similar, and therefore lie on the diagonal of the plot. In contrast, genes with spurious enrichment in one or two embryos lie below the diagonal. The gray line shows our exclusion criterion: 100 times difference between the largest and the third-largest number of cells with >10 UMI counts. The red dots mark 249 genes excluded for Figure 4e. Almost every embryo exhibited some of these spuriously enriched genes.

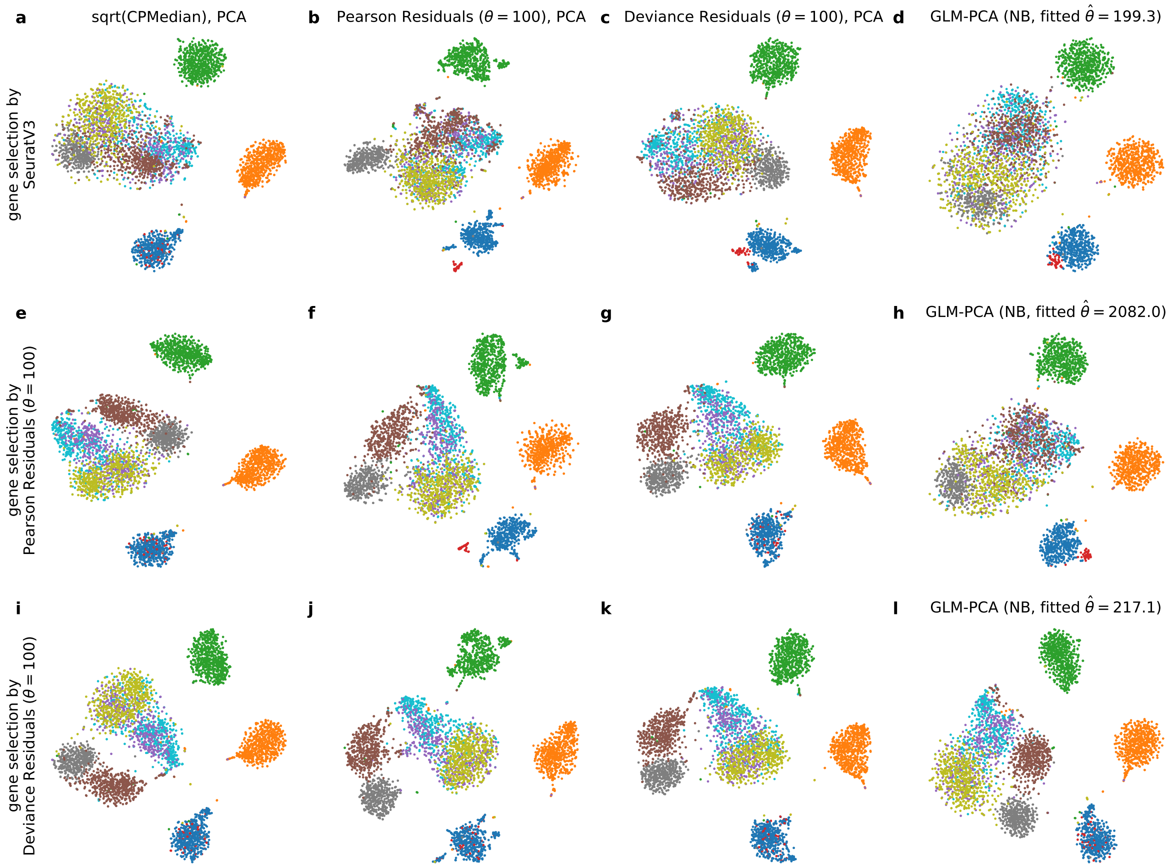


**Figure S7: t-SNE embeddings of the benchmarking dataset for selected normalization pipelines.** Rows show three HVG selection approaches, columns show four normalization and dimensionality reduction approaches. Each panel corresponds to one of the entries in Figure 5c.

### References

[1] Svensson V, Natarajan KN, Ly LH, Miragaia RJ, Labalette C, Macaulay IC, et al. Power analysis of single-cell RNA-sequencing experiments. Nature Methods. 2017;14(4):381–387.

[2] Klein AM, Mazutis L, Akartuna I, Tallapragada N, Veres A, Li V, et al. Droplet barcoding for single-cell transcriptomics applied to embryonic stem cells. Cell. 2015;161(5):1187–1201.

[3] Han X, Wang R, Zhou Y, Fei L, Sun H, Lai S, et al. Mapping the mouse cell atlas by microwell-seq. Cell. 2018;172(5):1091–1107.

[4] Grün D, Kester L, Van Oudenaarden A. Validation of noise models for single-cell transcriptomics. Nature methods. 2014;11(6):637–640.

[5] Hafemeister C, Satija R. Normalization and variance stabilization of single-cell RNA-seq data using regularized negative binomial regression. Genome Biology. 2019;20:296.

[6] Townes FW, Hicks SC, Aryee MJ, Irizarry RA. Feature selection and dimension reduction for single-cell RNA-Seq based on a multinomial model. Genome Biology. 2019;20:295.

[7] Macosko EZ, Basu A, Satija R, Nemesh J, Shekhar K, Goldman M, et al. Highly parallel genome-wide expression profiling of individual cells using nanoliter droplets. Cell. 2015;161(5):1202–1214.

[8] Wagner F. Straightforward clustering of single-cell RNA-Seq data with t-SNE and DBSCAN. BioRxiv. 2019;p. 770388.
